# PTPN22 interacts with EB1 to regulate T cell receptor signaling

**DOI:** 10.1101/481507

**Authors:** Xiaonan Zhang, Bin Bai, Tao Wang, Jiahui Zhao, Na Zhang, Yanjiao zhao, Xipeng Wang, Yang Yu, Bing Wang

## Abstract

PTPN22 has been reported as an important negative regulator of T cell signaling. Here we identified EB1 as an associated protein of PTPN22 via 2-hybrid and mass spectrometry screening.

Recently the phosphorylation of EB1 has been proved in the regulation of T cell receptor (TCR) mediated signaling pathway. Our results shown that PTPN22 interacted with EB1 through the P1 domain of PTPN22, and regulated the Y247 phosphorylation site of EB1. The subsequent results suggest that PTPN22 interacts with EB1 and regulate the phosphorylation of EB1, which results in the regulation of the expression of T cell activation markers of CD25 and CD69, and the phosphorylation levels of the T cell signaling molecules, such as ZAP-70, LAT and Erk, ultimately resulting in NFAT transcription factors entering the nucleus and regulating the secretion of cytokine IL-2. This newly identified interaction between PTPN22 and EB1 may play an important role in TCR signal pathways.

## Introduction

Protein tyrosine phosphatases (PTPs) are essential signal transduction enzymes that mediate the immune response [1]. Inheritance of a coding variant of the protein tyrosine phosphatase nonreceptor type 22 (PTPN22) gene is associated with increased susceptibility to autoimmunity and infection [2, 3]. The PTPN22 gene is located on the short arm of chromosome 1 (1p 13.3-lp l3.1) and encodes Lyp/Pep protein tyrosine phosphatase, which is mainly expressed in lymphocytes to inhibit the functions of T and B cells. The 802-residue of PTPN22 protein structure contains an N-terminal canonical phosphatase domain, and the five C-terminal proline, glutamic/aspartic acid, serine, threonine (PEST)-containing sequences [4]. During recent years, PTPN22 has been reported as a negative regulator of the TCR signaling pathway. For example, PTPN22 is involved in PTPN22-Csk interactions, which causes the activation of tyrosine kinase Csk to phosphorylate the inhibitory tyrosine sites on Lck and Fyn of the Src family tyrosine kinases, while four proline-rich regions of PTPN22 act as mediates by binding to the Src homology 3 (SH3) domain of the Csk tyrosine kinase [5-10]. The human PTPN22 gene encoding Lyp has been associated with a number of autoimmune diseases including rheumatoid arthritis [11-13], Type I diabetes [12, 14], Graves’ disease [15], and systemic lupus erythematosus [16].

End-binding proteins (EBs) are highly conserved and ubiquitous plus-end tracking proteins (+TIP) [17]. The N-terminal calponin homology (CH) domain of EB1, EB2 and EB3 is able to bind to MTs. The C-terminus of EBs have a flexible acidic tail that contains the sequence EEY/F, which is required for self-inhibition and binding to various partners, and can also control MT growth [18, 19]. End-binding protein 1 (EB1) is a member of microtubule +TIP, which bind to the plus ends of growing MTs via its conserved calponin-homology domain [20]. EB1 regulates MT stability at bundled actin-rich sites of the cell cortex through the recruitment of other +TIPs, such as SxIP motif-containing proteins and cytoskeleton-associated glycine-rich domain (CAP-Gly) containing proteins, to MT plus ends [21, 22] or by coordinating with other MT-binding proteins such as adenomatous polyposis coli (APC), CLASPs, MT-actin crosslinking factor 1 (MACF1)/actin crosslinking family 7 (ACF7), and mitotic centromere associated kinesin (MCAK) [23-25]. EB1 plays an important role in the regulation of microtubule dynamics and has been implicated in many autoimmune diseases [26, 27]. Post translational modification of EB1 especially through phosphorylation and dephosphorylation is very important for the normal function of EB1 proteins. Recently, EB1 was found to bind to the CD3 ITAMs and also regulate vesicular trafficking at the IS, and therefore is the connection between the TCR and downstream signaling molecules [28]. Ran et al. reported that the phosphorylation of EB1 regulates the recruitment of CLIP-170 and p150 to the plus ends of astral microtubules [29].

However, to date, there are no PTPs have been identified to associate with or regulate the phosphorylation levels of EB1. Herein, we identified EB1 as an associated protein of PTPN22 via 2-hybrid and mass spectrometry screening. Our results shown that PTPN22 interacted with EB1 through the P1 domain of PTPN22, and regulated the Y247 phosphorylation site of EB1, which results in the regulation of the expression of T cell activation markers of CD25 and CD69, and the phosphorylation levels of the T cell signaling molecules, such as ZAP-70, LAT and Erk, ultimately resulting in the secretion of cytokine IL-2. This newly identified interaction between PTPN22 and EB1 may play an important role in TCR signal pathways.

## RESULTS

### PTPN22 interacts with EB1 in vitro and in vivo

Virtually all processes in living cells are dependent on protein-protein interactions (PPIs). The most frequently used powerful genetic systems for mapping PPIs are the yeast two-hybrid (Y2H) screen and mass spectrometry analysis. In this study, yeast two-hybrid screens were performed to screen a mouse hematopoietic cells cDNA library [30]. The carboxy-terminal domain of PTPN22_601-802_ was employed as a bait to screen potential proteins which may interact with PTPN22. About 50 positive colonies had grown from one million transfectants, and the EB1 were one of the top candidates among the selected colonies (Figure 1A). Interestingly the EB1 was also identified as a binding partner of PTPN22 via mass spectrometry analysis (Figure 1B, EV1). To test the physical interaction between PTPN22 and EB1, we performed a GST pull-down assay *in vitro* (Figure 1C, EV2), and found that His-EB1 was pulled down with the immunoprecipitate of GST-PTPN22, suggesting the possibility that PTPN22 can interact with EB1 *in vitro*. In addition, Flag-PTPN22 was verified to associate with EB1 by a co-immunoprecipitation (CO-IP) assay of an overexpression system in 293T cells (Figure 1D), then the interaction between endogenous PTPN22 and EB1 was performed in BI-141, Jurket and RAW264.7 cell lines was studied with the use of CO-IP assays, with the results showing that endogenous PTPN22 and EB1 proteins may form a complex (Figure 1E, F, G). Subsequently, the interaction between PTPN22 (Green fluorescence protein) and EB1 (Red fluorescence protein) was also confirmed in Jurket cells and RAW264.7 cells through the cell co-localization assay (Figure 1H, I).

**Figure 1.**
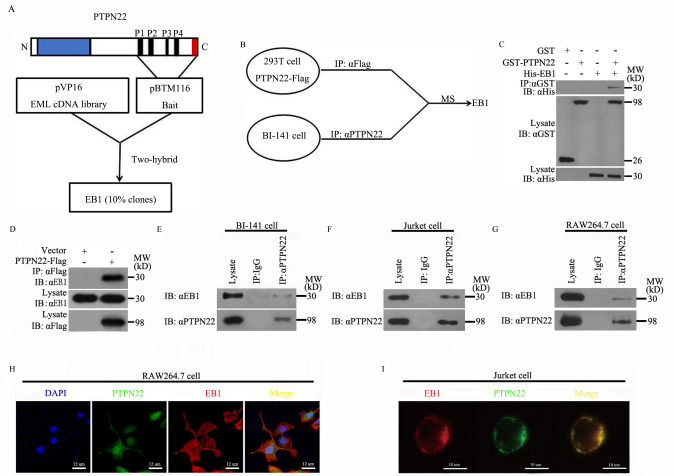
PTPN22 interacts with EB1 in vitro and in vivo. A Diagrammatic representation of the yeast two-hybrid system, which was performed to screen a mouse hematopoietic cells cDNA library constructed in pVp16 vectors, while the carboxy-terminal domain of PTPN22_601-802_ was employed as a bait to fuse to the Lex A, a DNA activation domain (AD) in the plasmid of pBTM116. As a result that 10% of the positive clones were EB1. B Affinity purification coupled to mass spectrometry mostly detects stable interactions and thus better indicates functional in vivo PPIs. The lysates of 293 T cells expressed with PTPN22-Flag were IP by anti-Flag antibody, while the lysates of BI141 T cells were IP by anti-PTPN22 antibody. PPIs can then be quantitatively and qualitatively analysed by mass spectrometry, and the results revealed that EB1 was colocalized in the PPIs. C GST pulled down assays indicate that recombinant GST-tagged PTPN22 but not GST can interact with His-tagged EB1. D Co-IP assay shows that tagged PTPN22 can interact with EB1 in 293T cell. E, F, G Co-IP assay shows that endogenous PTPN22 and EB1 can form a complex in Different cell lines. BI-141 cells (E) or Jurket cells (F) or RAW 264.7 cells (G). H, I Cell co-localization assays were used to confirmed the interaction between PTPN22 (Green fluorescence protein) and EB1 (Red fluorescence protein) in both cell lines. RAW 264.7 cells (H) or Jurket cells (I). Data information: Experiments were carried out twice (C-G) or three times (D) with similar results.

### Identifiion of the binding sites between PTPN22 and EB1

In order to identify the interaction sites between PTPN22 and EB1, the PTPN22 gene was divided into different fragments according to its the secondary protein structure of PTPN22 (PTPN22-Flag ΔC1, PTPN22-Flag ΔC2, PTPN22-Flag ΔC3, PTPN22-Flag ΔN1, PTPN22-Flag ΔN2 andPTPN22-Flag ΔN3) (Figure 2A). These fragments were transfected into 293T cells respectively. The association between Flag tagged PTPN22 fragments and endogenous EB1 were studied using CO-IP. As a results, PTPN22-Flag ΔC2, PTPN22-Flag ΔC3 and PTPN22-Flag ΔN1 can interact with EB1 (Figure 2C). Since these three fragments have a common P1 domain we constructed a P1 domain deleted PTPN22 mutant (PTPN22-Flag ΔP1) (Figure 2B). As shown in figure 2D PTPN22-Flag WT can interact with EB1 whereas PTPN22-Flag ΔP1 lost the binding features to EB1, which proving that P1 is the domain of PTPN22 to associate with EB1. Next the binding sites of EB1 need to be clarified. Figure 3A dipicts the structural model of EB1. It was reported that Tyr_217_ and Tyr_247_ of EB1 was the potential sites of the tyrosine phosphorylation, and that coiled-coil and EBH region are the site through which EB1 interacts with other molecules [31-33]. Here we constructed HA-EB1 Y217F, HA-EB1 Y217D, HA-EB1 Y247F, HA-EB1 Y247D to identify the binding sites of EB1 to PTPN22, and found that the HA-EB1 Y217F will not affect the interaction to PTPN22, while HA-EB1 Y247F/HA-EB1 Y247D decrease the binding to PTPN22 compared with EB1 WT. However, HA-EB1 Y217D completely lost the characteristics of interacting to PTPN22 (Figure 3B). This may be the mutant of Y to D on EB1 217 will changing the local conformation of EB1 protein to prevent the PTPN22/EB1 interaction [31].

**Figure 2.**
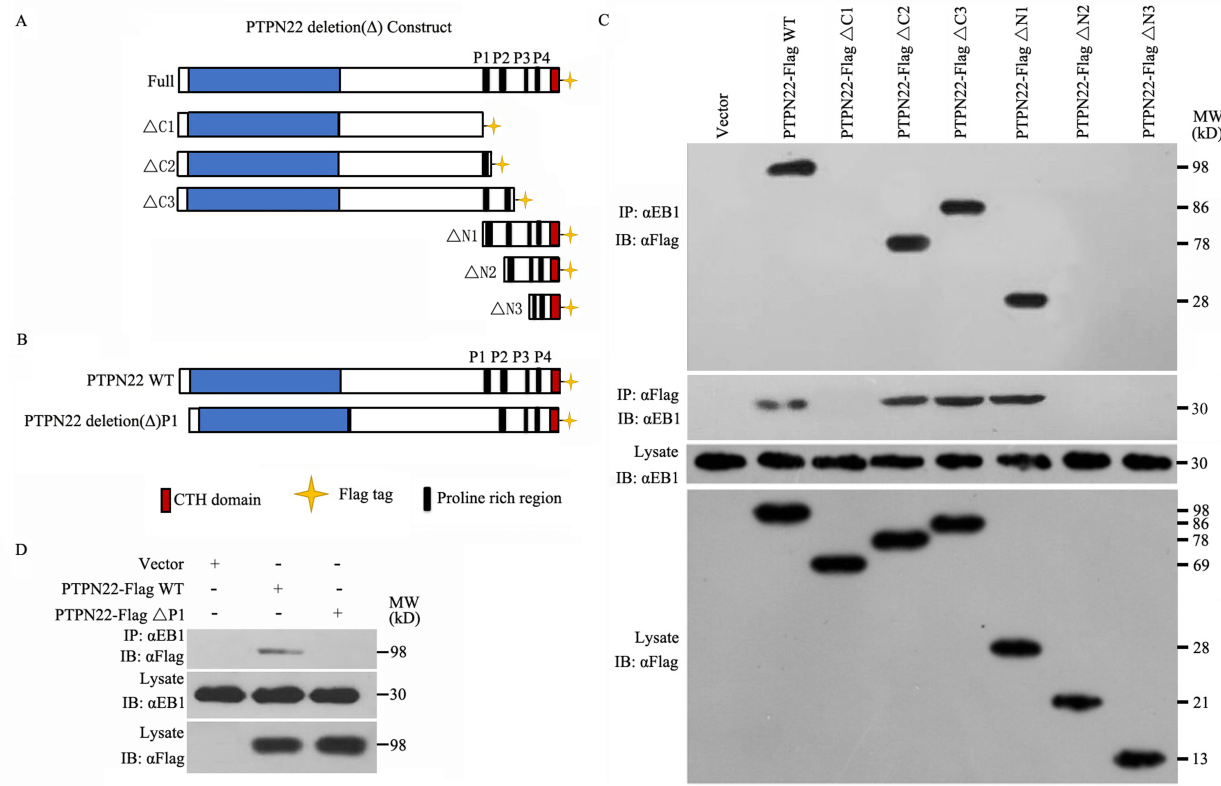
The P1 domain of PTPN22 is the domain that directly interacts with EB1. A Diagrammatic representation of the structures of different PTPN22 fragments. B Diagrammatic representation of the structure of PTPN22-Flag ΔP1. C Various truncated forms of PTPN22-Flag gene constructs were transfected in 293 T cells and the associations between the two proteins monitored by immunoblotting of anti-EB1 immuno-precipitates with antibodies against Flag (top panel), while the second panel indicated that the associations between the two proteins monitored by immunoblotting of anti-Flag immuno-precipitates with antibodies against EB1. The lower two panels show the parallel expressions of EB1 and the PTPN22 variants, respectively. D The Flag tagged P1 domain deleted mutant of PTPN22 (PTPN22-Flag ΔP1) was constructed, and using the CO-IP assay to verify that PTPN22-Flag ΔP1 cannot interact with EB1. Data information: Experiments were carried three times (C,D) with similar results.

**Figure 3.**
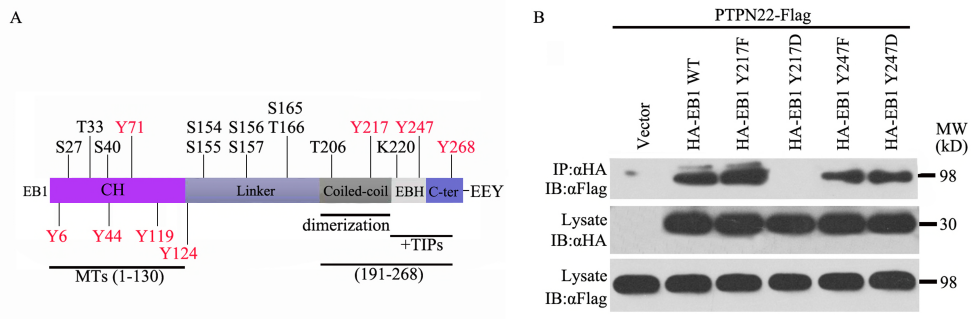
Tyr217 and Tyr247 of EB1 were the potential sites that were regulated by PTPN22. A Diagrammatic representation of the structure of EB1 protein. B CO-IP assay was performed between HA-EB1 Y217F, HA-EB1 Y217D, HA-EB1 Y247F, HA-EB1 Y247D and PTPN22-Flag, and the result shown that the HA-EB1 Y217D mutant cannot interact with PTPN22, whereas the other mutants of EB1 do bind to PTPN22. Data information: Experiments were carried three times (B) with similar results.

### PTPN22 dephosphorylated the 247 tyrosine site of EB1

PTPN22 acts as a tyrosine dephosphorylase to regulate the phosphorylation of physiological signaling pathways. CD3 and CD28 are commonly used to activate the phosphorylation of EB1. Therefore, we chose 15 min as an optimal stimulation condition and found that the phosphorylation levels of EB1 decreased significantly after transfection of PTPN22-Flag, indicating that EB1 could be a substrate of PTPN22 (figure 4A, B). Recent literatures have shown that Y247 is the primary residue in EB1 that is phosphorylated by Src kinase [31] while EB1 Y217D cannot dimerize or interact with other +TIPs, suggesting that phosphorylation of this residue may help maintain the equilibrium between the monomer and dimer forms of EB1, thereby regulating MT dynamics [33]. Thus, tyrosine sites of 217 and 247 were mutated and found that these two sites could be the phosphorylation sites since the phosphorylation levels of HA-EB1 Y217F, HA-EB1 Y247F were reduced compared with EB1WT (figure 4C shown), hence over expression of PTPN22 was utilized as shown in figure 4D that the phosphorylation levels of HA-EB1 Y217F decreased, whereas HA-EB1 Y247F did not change, indicating that Y247 was the site of EB1 that was regulated by PTPN22.

**Figure 4.**
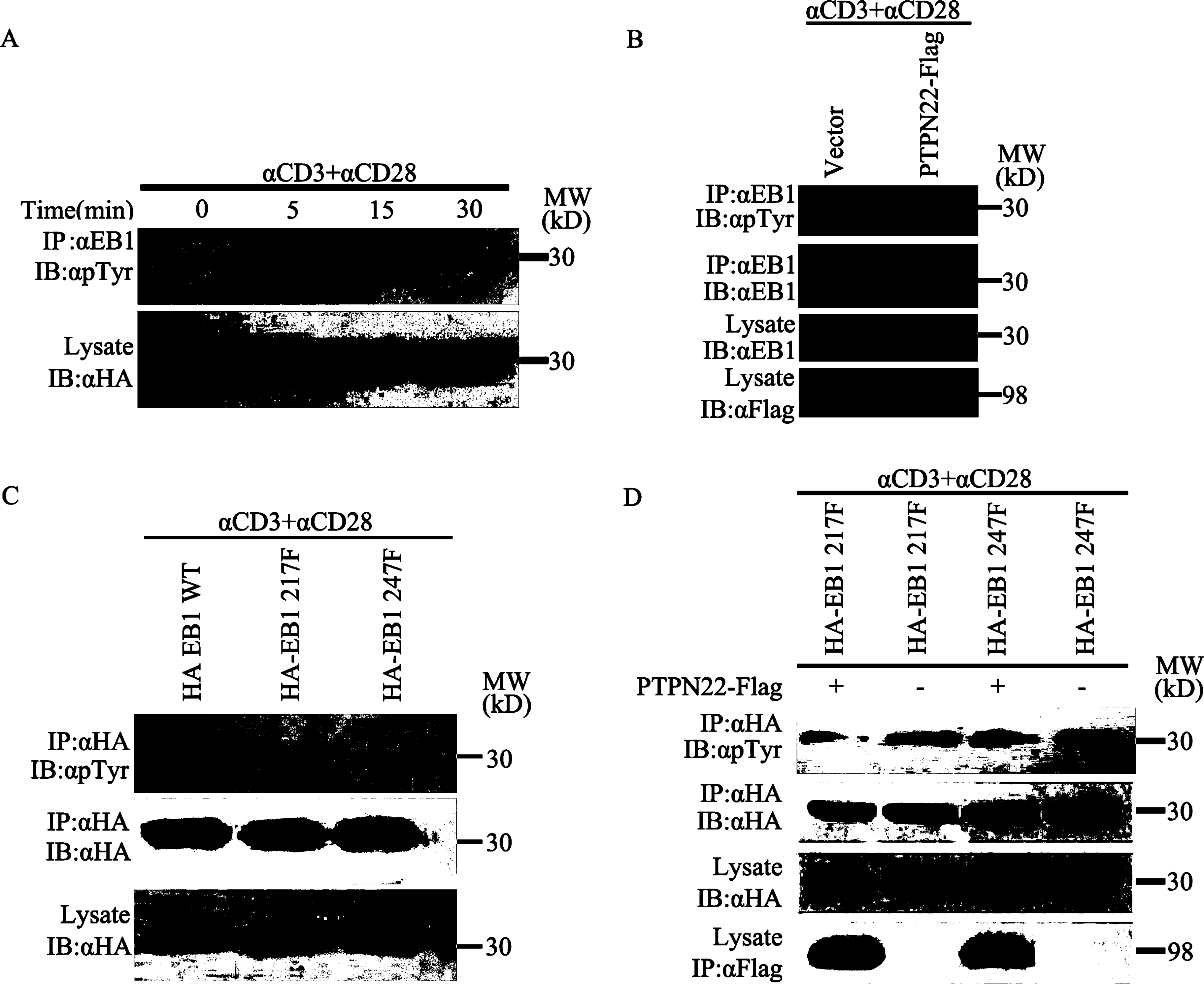
PTPN22 regulates the phosphorylation of EB1 through Tyr247. A The time course of TCR stimulation of BI-141 cells to monitor the phosphorylation levels of EB1, and 15min was selected as the optimal induction time for EB1 phosphorylation. B The phosphorylation levels of EB1 were significantly decreased after transfection of exogenous PTN22-Flag. C The phosphorylation levels of EB1 decreased significantly after transfection of exogenous HA-EB1 Y217F and HA-EB1 Y247F. D Co-transfection of PTPN22-Flag and HA-EB1 Y247F did not result in a significant change of the phosphorylation levels of EB1, however, co-transfection of PTPN22-Flag and HA-EB1 Y217Fdecreased the phosphorylation levels of EB1. Data information: Experiments were carried three times (A-D) with similar results.

### The binding domains between PTPN22 and EB1 are important on the activation of TCR signal pathways

It has been reported that PTPN22 is a negative regulator of TCR signal pathways, while the P1 domain of PTPN22 has been identified to be the site of interaction with EB1(figure 2D). In order to study the role of the P1 region in TCR signal pathways, the phosphorylation levels of immune activated upstream factors ZAP-70 and Lck, and downstream factors Erk were tested between PTPN22 ΔP1 and PTPN22 WT transfected cells. Compared with PTPN22 WT, PTPN22 ΔP1 up regulated the phosphorylation levels of ZAP-70, Lck and Erk remarkably (Figure 5A). CD25 and CD69 are early activation markers that are expressed in T cells and many other cell types in the immune system, while IL-2 is a cytokine secreted by the activated T cells. Western blot and qPCR were used to detect the protein and mRNA levels of the activation markers regulated by the P1 deleted mutant of PTPN22 ΔP1, as the results, PTPN22 ΔP1 can significant increase the protein and mRNA levels of CD25, CD69 and IL-2 (Figure 5B, C and D). Meanwhile, the enzyme activity of PTPN22 was not affected after the P1 domain was deleted (Figure 5E, F and G). The results implied that the deletion of the P1 domain blocked the interaction between PTPN22 and EB1, and eventurely reversed the inhibitory effect of PTPN22 on T cell immune activation without loss of enzyme activity.

**Figure 5.**
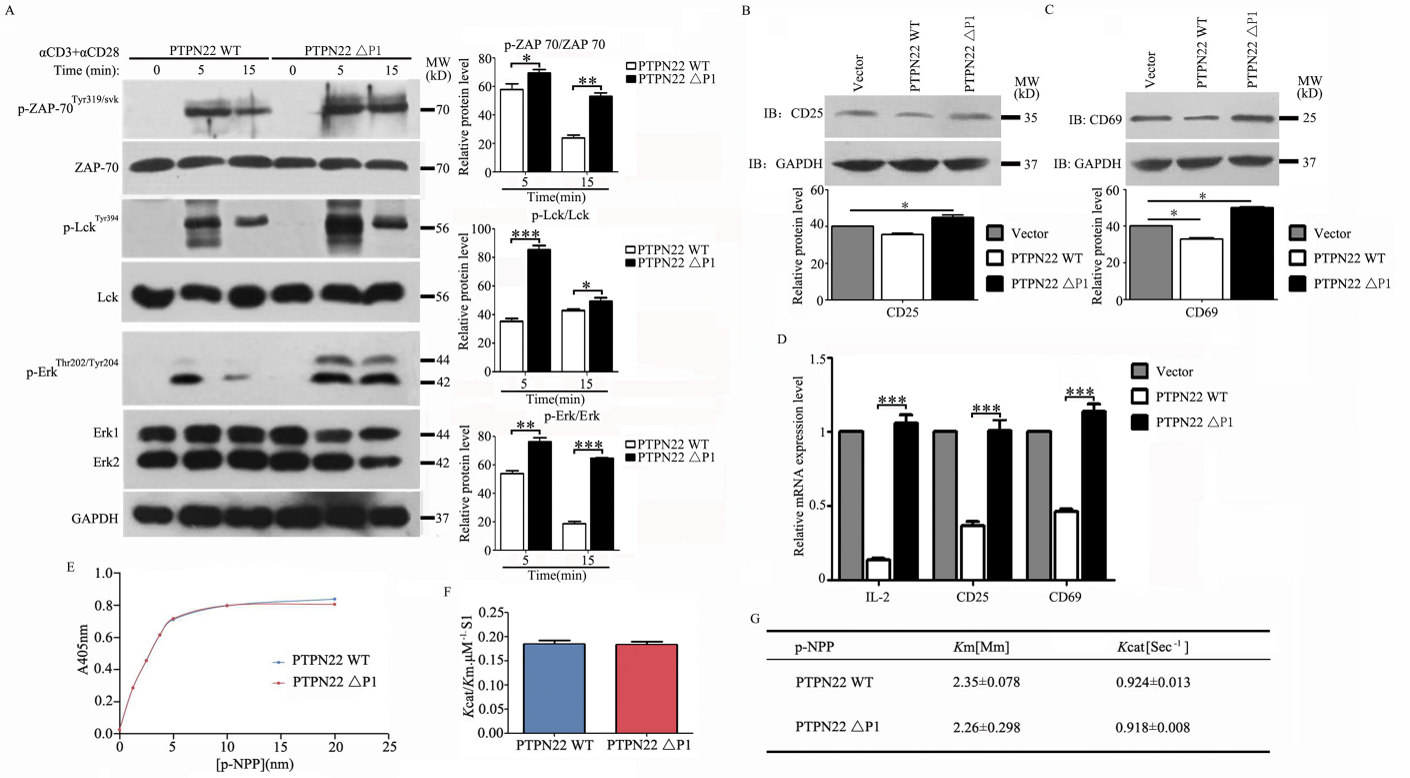
Deletion of the P1 domain of PTPN22 can reverse the inhibitory effect of PTPN22 on T cell immune activation. A Compared with the PTPN22 WT, PTPN22 ΔP1 can increase remarkably the phosphorylation levels of ZAP-70, Lck and Erk. B, C, D Western blot and qPCR assays were used to test the protein and mRNA expression levels of CD25, CD69 and IL-2, and the results shown that the protein and mRNA expression levels of CD25, CD69 and IL-2 increased significantly in PTPN22 ΔP1 transfected cells. Western blot assays were used to test the protein expression levels of CD25 and CD69 (B-C) or qPCR assays were used to test the mRNA expression levels of CD25, CD69 and IL-2 (D). E, F, G Deletion of P1 domain of PTPN22 will not affect its enzyme activity. pNPP hydrolysis with PTPN22 catalytic domain and PTPN22 ΔP1 (E) or Kinetics parameters for the wild-type and the mutants of PTPN22 toward pNPP (F,G). Data information: All Experiments were carried three times (A-E) with similar results. In (A-C), Significant differences were calculated with t-test and are indicated with *P < 0.05, **P < 0.01, ***P < 0.001.

Similarly, EB1 Y217D lost the feature of binding to PTPN22 comparied with EB1WT (figure 3B), therefore, EB1 Y217D transfected cells were used to study the phosphorylation of upstream immune response factors (ZAP-70, Lck, Fyn) and downstream immune response factors (Erk, Akt, p38), as shown in figure EV3A and figure 6. Compared with EB1 WT transfected cells, the phosphorylation levels of ZAP-70, Lck, Erk, Akt and p38 were up-regulated, while the phosphorylation level of Fyn were not affected. The results indicated that the interaction between PTPN22 and EB1 can affect the phosphorylation of immune response factors in TCR signal pathways.

**Figure 6.**
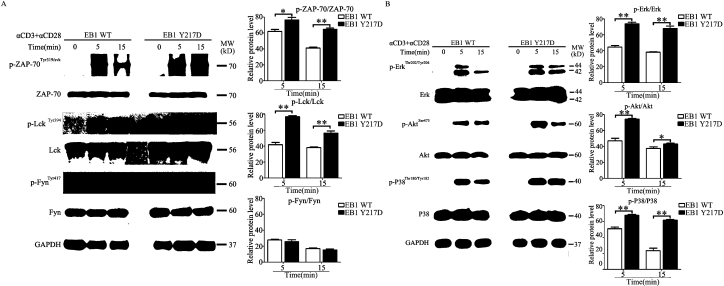
Interaction between PTPN22 and EB1 can regulate the phosphorylation of immune response factors. A The effects of EB1 Y217D on the phosphorylation levels of upstream immune response factors (ZAP-70, Lck, Fyn). B The effects of EB1 Y217D on the phosphorylation levels of downstream immune response factors (Erk, Akt, p38). Data information: Experiments were carried three times (A,B) with similar results. Significant differences were calculated with t-test and are indicated with *P < 0.05, **P < 0.01, ***P < 0.001.

### PTPN22 regulates the immune responses by regulating the phosphorylation of EB1

Mutation of C227 in PTPN22 to S227 can make PTPN22 lose the activity of dephosphorylase. We compared the effect of PTPN22 WT with PTPN22 C227S on the phosphorylation levels of CD3ζ, ZAP-70, LAT, PLCγ1, PKCδ and Erk. According to the results, the phosphorylation levels of CD3ζ, ZAP-70, LAT, PLCγ1, PKCδ and Erk in PTPN22 C227S transfected cells were significantly up regulated (Figure 7A, EV3C). Tyrosine 247 of EB1 was the site regulated by Src and PTPN22 (Zhang et al 2016, Figure 4D). Mutation of Y247 in EB1 to F247 blocked the phosphorylation of EB1, as shown in figure EV3B and 7B, compared with the wild type EB1 Y247F mutant decreased significantly the phosphorylation levels of CD3ζ, ZAP-70, LAT, PLCγ1, PKCδ and Erk in EB1 Y247F transfected cells. The results indicate that PTPN22 may regulate the phosphorylation of downstream immune activating factors by regulating the phosphorylation of EB1, which finally leads to the regulation of immune responses.

**Figure 7.**
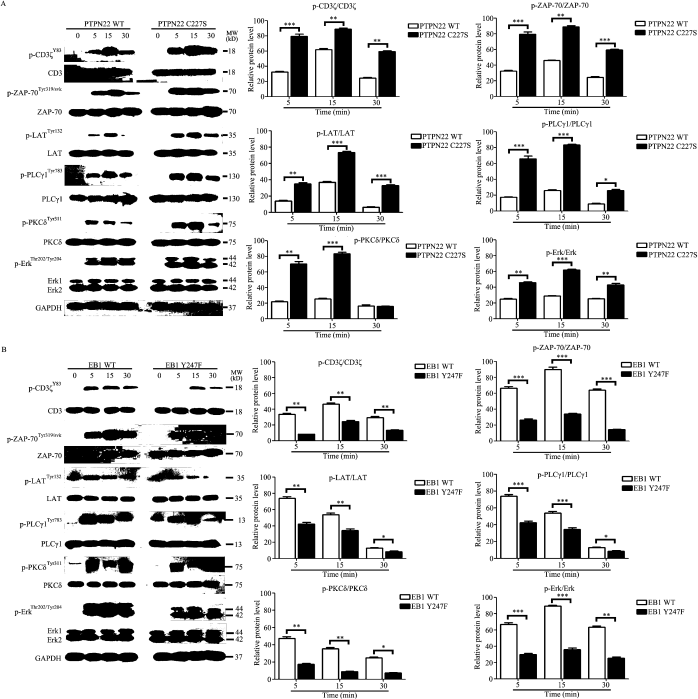
PTPN22 regulates the immune responses by regulating the phosphorylation of EB1. A The effects of PTPN22 WT and PTPN22 C227S on the phosphorylation levels of CD3ζ, ZAP-70, LAT, PLCγ1, PKCδ, and Erk were tested by Western blot assay, compared with PTPN22 WT transfected cells, the phosphorylation levels of PLCγ1, LAT, PKCδ in PTPN22 C227S transfected cells were up regulated significantly. B The effects of EB1 WT and EB1 Y247F on the phosphorylation levels of CD3ζ, ZAP-70, LAT, PLCγ1, PKCδ and Erk were tested by Western blot assay. Data information: Experiments were carried three times (A,B) with similar results. Significant differences were calculated with t-test and are indicated with *P < 0.05, **P < 0.01, ***P < 0.001.

### Mutation of the PTPN22 binding or dephosphorylation sites of EB1 can regulate the TCR signaling pathways

CD25 and CD69 are activation markers of T cells. PTPN22 ΔP1, a mutant is not able to associate with EB1, can significant increase the protein and mRNA levels of CD25, CD69 and IL-2 (Figure 5B, C and D). In order to further study the role of PTPN22/EB1 in TCR signalings, after the transfected BI-141 T cells were stimulated by SEB, CD3, CD3 + CD28, EB1 Y217D and EB1 Y247F, the binding and dephosphorylation sites of PTPN22, were utilized to assay the protein expression levels of CD25 and CD69 by enzyme-linked immunosorbent assay (ELISA) (Figure 8A and B), or the mRNA expression of CD25 and CD69 by qPCR (Figure 8C and D). According to the results, compared with the EB1 WT group, the expression of CD25 and CD69 increased significantly in EB1 Y217D transfected cells, whereas the expression of CD25 and CD69 decreased significantly in EB1 Y247F transfected cells. This implies that PTPN22 regulates the expression of CD25 and CD69 by regulating the phosphorylation of EB1.

**Figure 8.**
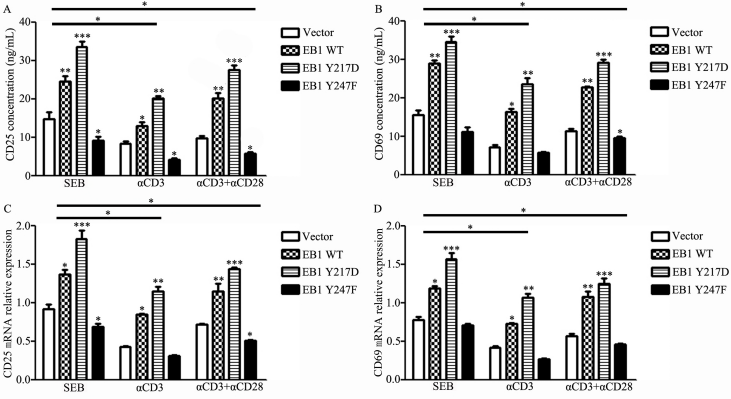
Phosphorylation of EB1 can regulate the expression of TCR signaling. A, B Test the protein expression levels of CD25 and CD69 by use of ELISA. ELISA assays were used to test the expression levels of CD25 (A) or Same assays were used to test the expression levels of CD69 (B). C, D Test the mRNA transcription levels of CD25 and CD69 by use of qPCR. qPCR assays were used to test the expression levels of CD25 (C) or Same assays were used to test the expression levels of CD69 (D). Data information: Significant differences were calculated with t-test and are indicated with *P < 0.05, **P < 0.01, ***P < 0.001.

Likewise, the downstream activation factors of TCR signal pathways were investigated as shown in figure 9. The Vector, EB1 WT, EB1 Y217D and Y247F were transferred into cells, and the cells were stimulated by SEB, CD3, CD3+CD28 for 30 min, then dual luciferase reporter assay was used to test the transcription factor of NFAT and secretion of IL-2. The results revealed that compared with EB1 WT group, the transcription factor of NFAT and secretion of IL-2 increased significantly in EB1 Y217D group and decreased significantly in EB1 Y247F transfected cells (Figure 9A, B), while the mRNA expression of NFAT and IL-2 by qPCR (Figure 9C, D). These results implied that the association sites of both PTPN22 and EB1 or the phosphorylation of EB1 maybe invovled in the T cell activation by TCR signal pathways.

**Figure 9.**
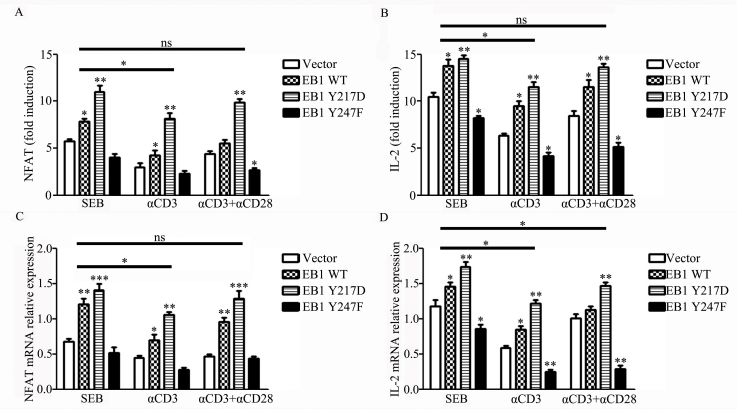
phosphorylation of EB1 regulates NFAT transcription factors and secretion of IL-2. A, B Dual luciferase reporter assay was used to determine the role of phosphorylation of EB1 on T-cell activation transcription factor of NFAT and IL-2. Dual luciferase reporter assays were used to test the expression levels of NFAT (A) or Same assays were used to test the expression levels of IL-2 (B). C, D The mRNA expression levels of NFAT and IL-2 were monitored by using qPCR. qPCR assays were used to test the expression levels of NFAT (C) or Same assays were used to test the expression levels of IL-2 (D). Data information: Significant differences were calculated with t-test and are indicated with *P < 0.05, **P < 0.01, ***P < 0.001.

## Discussion

In this study, we reported that PTPN22, an immune response negative regulatory protein, can interact with the microtubule positive end binding protein EB1 *in vivo* and *in vitro*. The P1 domain of PTPN22 was the region that interacted with EB1, while Y247 was the site of EB1 that was regulated by PTPN22. We also identified that PTPN22/EB1 interaction may play an important role in regulating TCR signal pathways and EB1 is an important TCR signaling mediator, which is newly identified function of EB1.

Previous studies have reported that PTPN22 is a negative regulator of immune signaling, and that its structure is characterized by a phosphatase domain at the N-terminus and by four C-terminal PEST domains (defined as proline, glutamic/aspartic acid, serine, threonine) named P1-P4 which was a region rich in proline [4]. These PEST domains mediate protein-protein binding interactions. For expamle, CSK interacts with P1 domain of PTPN22[34, 35]. In this study through yeast two hybrid and MS assays, we identified that EB1 may be a new potential substrate for PTPN22, and PTPN22/EB1 interaction was confirmed *in vivo* and *in vitro* through subsequent co immunoprecipitation and pull down experiments. In order to find the specific interaction sites of PTPN22/EB1 interaction, we constructed a P1 deletion PTPN22 mutant (PTPN22 ΔP1), and the results showed that PTPN22 ΔP1 cannot binding to EB1, which indicates that the P1 domain of PTPN22 is involved in its association with EB1. Literatures revealed that the catalytic phosphatase domain of PTPN22 were important in regulation of the phosphorylation levels of LCK, Erk, Akt, p38, PLC, PKC, LAT, ZAP-70, and Fyn et al when T cells were activated via TCR pathways [7, 36]. In this study, when the PTPN22 ΔP1 overexpressed T cell was activated by TCR cell signaling pathways, the phosphorylation levels of tyrosine kinase ZAP70 and LCK were significantly up-regulated, as well as the phosphorylation levels of ERK signaling pathways. Further studies clarified that these results from the deletion of P1 were not caused through a change in PTPN22 enzyme activity, but is more likely due to the interaction with EB1. After being stimulated by specific antigens and mediated by TCR/CD3, there are many physiological changes induced in T cells, such as cell proliferation, apoptosis, differentiation, secretion of related cytokines, expression of the cytokine receptor etc. The changing of surface molecules such as CD25 and CD69 expression and the cytokine IL-2 secretion that are involved in the activation of T cells [37, 38]. Similarly, overexpression of PTPN22 ΔP1, a mutant cannot bind to EB1, increased the expression levels of T cell activation factors.

This lays a good foundation for us to further study the regulatory functions of the PTPN22/EB1 interaction in T cells. Therefore, we got to identify the binding domains of EB1 to PTPN22 and clarify the dephosphorylation sites of EB1 by PTPN22. The formation of two deep hydrophobic grooves on EBH mediates the interaction between EB protein and other + TIP proteins, and are also important domains for interaction with other various partners [18, 39]. For example, the N-terminal of CAP-GLY protein contains a CAP-GLY domain which directly binds to the EEY/F sequence at C-terminal of EB1, and mediates itself at the positive end of the growth microtubule; The C-terminal of CLIP170 also contains EEY/F sequences, which can bind to its own CAP-GLY structure and inhibit its interaction with EB1 [31]. Hence, the two tyrosine sites of EB1, Y217 and Y247 were mutated to D217 or F217 and D247 or F247, respectively (named as EB1 Y217D, EB1 Y217F, EB1 Y247D and EB1 Y247F, respectively). Immunoprecipitation assay was performed between these EB1 mutants and PTPN22. As a result, in addition to EB1 Y217D, other mutations can interact with PTPN22. It is reported that because the hydrophobic region of EBH can form a hydrogen bond with P or IP residues of SxIP, the EBH region of EB1 can interact with the SxIP domain of many proteins. But when Y217 is replaced by F or D, hydrogen bonds are opened, which may explain why Y217F decreases the interaction of MCAK and APC [40]. This involves changes in the three-dimensional structure, and suggests that EB1 Y217D does not interact with other proteins, and phosphorylation of this site may help maintain the balance between the EB1 monomer and dimer, thereby regulating the dynamics and normal function of microtubules.

Y247 is located in the EBH domain and forms two hydrogen bonds with E225 and T249. When Y247 is mutated into F or D, the hydrogen bond with E225 is abolished. This may change the stability of the hydrophobic region and affect the interaction between EB1 and SxIP domains of other microtubule positive-end interacting proteins. Therefore, the EB1 Y247F is not only a non-phosphorylated mutant, but this may also help explain why both PTPN22/EB1 Y247F and PTPN22/EB1 Y247D interaction were weakened. This is similar to the result reported by Yijun that EB1 Y247F and EB1 Y247D also weaken its combination with APC-C and MCAK [31]. In this study, the tyrosine phosphorylation of EB1 decreased in PTPN22 overexpressed cells, which preliminarily proved that PTPN22 could regulate the phosphorylation of EB1. In order to further identify specific regulatory sites of EB1, EB1 Y217F and EB1 Y247F overexpressed BI-141 cells were stimulated by CD3 and CD28, the phosphorylation levels of both EB1 Y217F and EB1 Y247Fdecreased, In order to prove whether PTPN22 had any effect on the phosphorylation of Y217 and Y247, PTPN22 overexpressed BI-141 cells were stimulated by CD3 and CD28, and compared with the control group. The phosphorylation of Y247F resulted in no obvious change, but the phosphorylation of Y217F decreased significantly in PTPN22 overexpressed cells, which proved that PTPN22 mainly regulated the tyrosine phosphorylation of Y247 of EB1in mouse T cells.

PTPN22 negatively regulates the signaling pathways of T cells. In resting T cells, Csk is anchored to the PAG/Cbp and binds to PTPN22 [41]. Phosphorylated PAG interacts with the SH2 domain of Csk, which in turn interacts with PTPN22, and eventually the complex inhibit the Src kinase family such as Lck that near cell membrane side [42]. When antigens activates TCR, CD45 dephosphorylates PAG/Cbp, resulting in Csk and PTN22 were released from the complex and separated from membrane [43]. Since the functions of PTPN22 ΔP1 on TCR pathways maybe due to the dissociation with Csk we investigated the effects of EB1Y217D, a mutant cannot bind to PTPN22, on the TCR pathways. The results revealed that, similar to PTPN22 ΔP1, EB1Y217D increased the phosphorylation levels of Lck, ZAP-70, Erk1/2, AKT and P38 compared to EB1 WT overexpressed cells. In addition, EB1 Y217D also increased the expression levels of CD69 and CD25 molecules, and the secretion of transcription factors NFAT and IL-2. Overall, these results also preliminarily demonstrate that PTPN22/EB1 interaction can affect the activation of TCR signaling pathways.

EB1 binds to ITAMs of CD3, and participates in the binding of T cells to APC cells through surface adhesion molecules and their receptors, which leads T cells from the state of motion to a highly polarized static state, and localized in the lymph nodes [27]. Our results have shown that PTPN22, which negatively regulates the T cell receptor signaling pathway, interacts with EB1 and regulates the phosphorylation Y247. After stimulation with specific antigens, and mediated by TCR/CD3, the ITAM motifs in the CD3 intracellular domain were phosphorylated by Lck. The phosphorylated ITAMs further recruited the intracellular tyrosine kinase ZAP-70. ZAP-70 can bind to a variety of junctional molecules, such as LAT, through its phosphorylated SH2 region, and cause the phosphorylation of Tyr in junctional molecules. Phosphorylated LAT can also bind to PLCγ, PKCδ and PI3K, then activate the Ras-Map kinase, the calcium-calcium phosphatase and, pKC pathways [44]. We found that EB1 Y247F decreased the phosphorylation levels of LAT, PLCγ1 and PKCδ compared with those in EB1 WT overexpressed cells. This suggests that the downstream signal of EB1 protein is involved in T cell activation that may be related to these molecules. EB1 has also been reported to control vesicle transport to immune synapses and can therefore regulate the association between TCR and downstream signaling molecules, such as LAT/PLCγ1 [44], These results suggest that PTPN22 may also affect vesicle transport and immune synapse formation by regulating EB1, but the specific mechanism remains to be studied.

EB1 Y247F can not only can lead to a change in EB1 phosphorylation levels, but can also led to structural changes that result in the loss of the hydrogen bond with E225. In order to confirm that the above results are likely to be caused by the regulation of PTPN22, the PTPN22 WT and C mutants PTPN22 C227S overexpressed T cells, were stimulated by CD3 and CD28. The resulting phosphorylation of ZAP-70,CD3ζ, LAT,PLCγ1,PKCδ and Erk in PTPN22 C227S overexpressed T cells were significantly higher than that of PTPN22 WT overexpressed cells, which further indicates that PTPN22 participates in the regulation of these downstream molecules, and indirectly indicates that EB1 Y247F regulates the phosphorylation of these molecules through dephosphorylation that weakens the activation of T cells.

As before, compared with EB1 WT group EB1 Y247F decreased the expression levels of CD25 and CD69, which further proves that PTPN22 regulates the activation of T cells by affecting the phosphorylation of ZAP-70,CD3ζ, LAT, PLCγ1, PKCδ and Erk and downstream transcription factors. Briefly, we found the direct interaction between PTPN22 and EB1, and that the P1 domain of PTPN22 was the key part of PTPN22/EB1 interaction, while it was also found that PTPN22 affects the phosphorylation of EB1 at Y247. Subsequent experimental results indicate that PTPN22 affects the EB1 phosphorylation, which in turn regulate the TCR signaling, while the elimination of EB1 dephosphorization by PTPN22 can initiate an immune response and increase the expression of immune response secretions such as IL-2, In conclusion, EB1 interacts with PTPN22 and acts as an important mediator for T cell receptor signaling. The study may provide a theoretical basis for the development of Y247 as a treatment target of T cell activation related autoimmune diseases.

## Materials and methods

### Antibodies and Chemicals

The antibodies used in this paper are as follows: mouse anti-Flag (F3165, Sigma-Aldrich), anti-HA (ab18181, Abcam), anti-GST (ab92, Abcam), anti-PTPN22 (#14693, CST), anti-EB1(#2164, CST), anti-His (ab9108, Abcam), anti-GAPDH (ab8245, Abcam), anti-pTyr (05-777, Merck millipore), anti-CD25 (ab156285, Abcam), anti-CD69 (ab202909, Abcam), anti-PLCγ1(#5690, CST), anti-LAT (ab2507, Abcam), anti-pLAT(Tyr132) (ab4476, Abcam), anti-ZAP-70 (ab32429, Abcam), anti-pZAP-70(Tyr319)(ab131270, Abcam), anti-Lck (ab3885, Abcam), anti-AKT (#4691, CST), anti-pAKT (Ser473) (#4060, CST), anti-P38 (#8690, CST), anti-pP38 (Thr180/Tyr311) (#4511, CST), anti-pLck(Tyr394) (ab201567, Abcam), anti-Fyn (ab1881, Abcam), anti-pFyn(Tyr417)(CST, #2101) anti-PKCδ (ab182126, Abcam), anti-pPKCδ (Tyr311) (ab76181, Abcam), anti-Erk (ab17942, Abcam), Anti-pErk1 (Thr202)/Erk2 (Tyr204) (#4370, CST), anti-pPLCγ1(Tyr783)(#2821, CST), anti-CD3ζ (ab226475, Abcam), anti-pCD3ζ (Tyr83) (ab68236, Abcam), Staphylococcal Enterotoxin B Domain (SEB) (144-153, Absin), SYBR Green qPCR Master Mix (HY-K0501, MedChem Express), Dual-Luciferase Report kit (E1910, Promega), 4’,6-diamidino-2-pheny-lindol (DAPI) (F3165, Sigma-Aldrich) and Protease inhibitor cocktail (Amresco, Solon, OH, USA).

### Yeast two-hybrid screening for protein interaction

During a yeast two-hybrid screen using LexA as part of the fusion proteins with a bait in the presence of the Src kinase. A partial PTPN22 cDNA (The cDNA encoded the carboxy-terminal domain of PTPN22) was cloned and inserted into the expression vector pBTM116/Src containing the SaII enzyme cut site. In addition to the DNA binding domain of LexA, the vector also had Src kinase (with tyrosine-to-phenylalanine mutations at positions 416 and 527) that could induce tyrosine phosphorylation of yeast cells. Correct expression of the bait was verified using western blotting of cell extracts using a mouse monoclonal antibody directed against the LexA domain (Santa Cruz Biotechnology, Inc., Dallas, TX). The absence of self-activation was verified by co-transformation of the bait together with a control prey and selection on a minimal medium lacking the amino acids tryptophan, leucine, and histidine (selective medium). For the yeast two-hybrid screens, the mouse hematopoietic cell (EML) cDNA Library of which expression vector VP16 contains a trans activator was transformed into yeast cells. Positive transformants were tested for β-galactosidase activity using a filter assay [45]. The identity of positive interactors was determined by DNA sequencing.

### Cell culture and Transfections

HEK 293T cells (ATCC, Rockville, MD) and RAW 264.7 cells (ATCC, Rockville, MD) were cultured in Dulbecco’s modified Eagle,s medium (Hyclone, Thermofisher) supplemented with 10% fetal bovine serum (Hyclone, Thermofisher). Antigen-specific mouse T-cell hybridoma BI-141 cell was a gift from Dr. André Veillette (IRCM), Jurkat T cell Clone E6-1 (ATCC, Rockville, MD) were cultured in RPMI 1640 (Hyclone, Thermofisher) supplemented with 10% FBS (Hyclone, Thermofisher), penicillin (1000U/ml) (Gibco, Invitrogen), and streptomycin (1000U/ml) (Gibco, Invitrogen). All the cells used here were cultured in a 37°C incubator with 5% CO_2_. Cells were transfected through the use of lipofectamine 3000 (Life technology, Thermo fisher) and DMRIE-C (Invitrogen), according to the manufacturer’s protocol and used after 24-48 hours.

### Protein expression, purification, and GST pull-down assay

The pGEX-4T-1-PTPN22, pGEX-4T-1 and pET-28a-EB1 were transformed in Escherichia coli strain BL21, respectively, and the GST, GST-PTPN22 or His-EB1 proteins were induced with 0.3 mM isopropyl-β-D-thiogalactopyranoside (IPTG). The cells were harvested after 20 hours of induction at 16°C, and were re-suspended in pre-cooled PBS, and homogenized by sonication. The lysates were cleared by centrifugation, and the supernatant was subjected to a GST tag protein purification gel and His tag protein purification gel (GE Healthcare) for affinity purification. GST and GST-PTPN22 proteins were eluted from the column using a 20 mM glutathione elution buffer, while His-EB1 protein was eluted from the column using 200, 250, and 400 mM imidazole elution buffer.

### GST Pull-Down Assays with EB Proteins

For the GST pull-down assays, equal amounts of purified GST or GST-PTPN22 proteins were bound to glutathione agarose (Fisher Scientific), according to the manufacturer’s instructions and the beads were washed four times using PBS. The recombinant His-tagged EB1 protein was then incubated with pull-down lysis buffer for 2 hours at 4°C. The eluted proteins were detected using SDS-PAGE and Western blot.

### Co-immunoprecipitation assay

The cells were harvested and lysed in pre-cooled Co-IP lysis buffer for 24-48 hours after transfection. The cell lysates were precipitated with a proposed mAb-conjugated Protein A sepharase (GE Healthcare, Fairfield, CT) or anti Flag magnetic beads and anti HA magnetic beads overnight at 4°C with gentle rotation. The beads were washed 4 times with 1 ml of the lysis buffer and then boiled with 1×SDS loading buffer for 5 minutes. The supernatants were subjected to SDS-PAGE and Western blot analysis.

### Enzyme Kinetics

PTPN22 and PTPN22 ΔP1 enzyme activity was tested according to a previously reported method. [46, 47]. All assays were performed at 37°C in 50 mM 3,3-dimethylglutarate (pH 7.0) buffer except for the experiment of pH dependency. 1 mM EDTA and 1 mM DTT were included in the 50 mM 3, 3-dimethylglutarate buffer, and the ionic strength was adjusted to 0.15 M using NaCl. Previous enzymological studies suggest that the PTPs catalysis have two steps [48, 49]. In the first step, ArOPO_3_ is the substrate. When pNPP was used as a substrate, the reaction was stopped by the addition of 1.0 M NaOH, and the activity was detected by monitoring the absorbance of paranitrophenol at 405 nm. All buffers contained 1 mM EDTA and 1 mM DTT, and were adjusted to an ionic strength of 0.15 M with NaCl. The Kcat value for pNPP hydrolysis was catalyzed by the wild type Lyp and was determined at 37°C. The following equation was used to fit the Kcat value against pH Kcat=(Kcat)^max^/((1+H/K_ES1_+K_ES2_/H)). In this equation, KES1 and KES2 are the apparent ionization constants of the enzyme-substrate complex in the rate-limiting step, and H is the proton concentration [50].

### ELISA

CD25 ELISA was performed using the pre-packaged CD25 ELISA kit (DY2438, R & D system), while CD69 ELISA was performed using the pre-packaged CD69 ELISA kit (TWp022633, www.tw-reagent.com) following the manufacturer’s protocols. All ELISA measurements were performed in triplicates.

### Cell Co localization Assay and Imaging

For the co-localization analysis of PTPN22 with the EB1, Raw 264.7 and Jurkat cells were grown on cover slips in 6-well plates at 70-80% confluence. The cells grown on glass coverslips were fixed with 4% paraformaldehyde and 0.12 mM sucrose in PHEM (60 mM PIPES, 25 mM Hepes, 5 mM EGTA and 2 mM MgCl_2_), and permeabilized for 5 minutes at room temperature with 0.2% Triton X-100 in immunfluorescence solution (PHEM containing 3% BSA, 100 ug/ml γ-globulin and 0.2% azide). The Cells were blocked for 30 minutes with an immunofluorescence solution and stained with the indicated PTPN22 and EB1 antibodies overnight at 4°C, and washed with PBS for three times, and were then incubated with secondary antibody for 1hour at 37°C. Nuclear DNA was then stained with DAPI for 4 minutes and images were obtained using a Nikon TE-2000E or Olympus confocal microscope.

### Dual-luciferase reporter Assay

The enzymatic activity of reporter proteins were quantified by using the Dual-Luciferase Reporter Assay System (Promega, Madison, WI, USA) according to the manufacturer’s instructions. After 48 hours of transfection with appropriate expression vectors that contained the promoter of IL-2 and NFAT, BI-141 cells were collected by centrifugation, lysed, and mixed with a substrate solution containing beetle luciferin, ATP, and either magnesium for firefly luciferase or coelenterazine for Renilla luciferase, and light emission was quantified, as described previously (Takahashi et al. 2009). The light emission background was determined with the vector, EB1 WT, EB1 Y217D and EB1 Y247F transfected cells. Each experiment was performed in triplicate [51].

### Quantitative RT-PCR assay

TRIzol reagent (Fisher Scientific) was used to extract the viral RNA from cellular samples. The procedure for quantitative RT-PCR was carried out as described previously [52]. Total RNA was isolated using TRIzol reagent (Thermo Fisher Scientific) according to the manufacturer’s instructions. The cDNA was reverse-transcribed from 1 μg of total RNAs using a Quant One Step RT-PCR Kit (TIANGEN, Beijing, China). 2μg of total RNA was converted to cDNA and the relative mRNA levels were quantified using a CFX96 Touch™ Real-Time PCR Detection System (Bio-Rad Laboratories). SYBR green chemistry was used. The primers are shown in Table EV2.

### Statistical analysis

The data is presented as means ± standard deviations (SD). The Graphpad Prism software (version 5.0) was used to determine the significance of the variability among different groups using two-way ANOVA test of variance. A p value< 0.05 was considered to be statistically significant.

## Acknowledgements

We thank Dr. Andre Veillette at IRCM for help with useful ideas and experimental materials. This research was supported by the National Natural Science Foundation of China (31670770, 2016YFC1302402, 31370784) and the Fundamental Research Funds for the Central Universities of China (N141008001, N162004006, N172008008).

## Author contributions

XZ performed the majority of the experiments, analyzed the data, wrote and edited the manuscript. BB and TW constructed of plasmids. JZ, NZ, YZ and XW performed ELISA, Dual-luciferase reporter Assay experiments and data analysis. BW and YY directed the study, analyzed and approved all of the data, wrote and edited the manuscript. All authors reviewed the manuscript.

